# Sex-Dependent Modulation of Emotional and Cognitive Processes by Prefrontal CB1 Receptors

**DOI:** 10.64898/2026.07.23.740323

**Authors:** M Ceprian, J Egaña-Huguet, L Godoy, A Godoy, A Aranguren-Alberdi, P Ospital, P Reyes-Velasquez, S Calovi, JA Santos-Martín, J Piriz, A Ramos-Miguel, S Mato, E Soria-Gomez

**Affiliations:** University of the Basque Country, EHU; Achucarro Basque Center for Neuroscience, Leioa Spain; Translational Neuropsychiatry Unit, Aarhus University, Aarhus, Denmark; Pharmacology Department, Sao Paulo University, Riberaio Preto, Brasil; Neuronal Excitability Lab, Instituto Biofisika (CSIC-EHU), Leioa, Spain; Ikerbasque, Basque Foundation for Science; Biobizkaia Health Research Institute, Baracaldo, Spain

## Abstract

The medial Prefrontal Cortex (mPFC) participates in emotional regulation, decision-making and behavioural flexibility. Cannabinoid receptor 1 (CB_1_) is widely expressed in the mPFC, particularly in GABAergic neurons, where it modulates synaptic transmission, contributing to the mPFC excitation-inhibition balance. Alteration of GABAergic activity and CB_1_ levels is indeed part of the pathophysiology of many psychiatric disorders, including depression, anxiety, and schizophrenia. Interestingly, both CB_1_ and mood disorders display important sex differences.

In this work, we study the **role of CB_1_ receptors in prefrontal GABAergic interneurons in emotional and cognitive processes in a sex-dependent manner**. To achieve this objective, we deleted CB_1_ from all mPFC neurons and the GABAergic population in adult CB_1_-flox male and female mice, and GABAergic neuronal activity was assessed via calcium imaging with fiber photometry.

Global CB_1_ deletion in mPFC neurons, specifically in GABAergic cells, altered **emotional** but not cognitive processes, with **opposite patterns**. This impairment was **sex- and task-dependent**. While pan-neuronal CB_1_ deletion had an anxiolytic effect on females, GABAergic CB_1_ deletion had the same effect on male mice, linked to increased GABAergic neuronal activity. By contrast, fear conditioning was primarily affected in males with neuronal CB_1_ depletion and in females with receptor deletion in inhibitory neurons. GABAergic CB_1_ deletion potentiates females’ freezing response during acquisition and recall 24 hours later, and is associated with decreased inhibitory neuronal activity during the tone-shock association.

In conclusion, mPFC GABAergic CB_1_ deletion is associated with an anxiolytic phenotype but also heightened responses to conditioned cues in a sex-dependent manner.

## Introduction

The Prefrontal Cortex (PFC) is a brain region involved in high-level tasks, including decision-making and emotional modulation^1–3^. To successfully perform these tasks, this region serves as a hub, receiving and processing information from different brain regions. In particular, the medial PFC (mPFC) has been implicated in fear recall and anxiety modulation, with key inputs from the hippocampus or the ventral tegmental area and outputs to the hypothalamus or amygdala^4^. GABAergic interneurons of the mPFC are key cellular substrates that integrate such inputs and outputs, thereby participating in the execution of complex behaviours^5^. In particular, mPFC-GABA alterations are described in several psychiatric disorders, such as schizophrenia or depression^6,7^.

Cannabinoid receptor 1 (CB_1_) is widely expressed in mPFC, particularly in GABAergic neurons^8,9^. Localised mostly at the presynaptic level and coupled to a Gi protein, its activation leads to neuronal inhibition^10,11^. Interestingly, a decrease in the CB_1_ receptor expression or changes in its binding properties have been described in the PFC of patients with schizophrenia^12,13^ or suicide patients^14^, where it may be linked to an Excitation / Inhibition imbalance. In this line, prenatal CB_1_ downregulation in mice leads to adult emotional and social impairment^15^.

Both PFC activity and CB_1_ affinity have a clear sex dimorphism^16–26^. In both rodents and humans, PFC activity patterns differ between sexes^17,18^, with differences in GABAergic signalling and tone^16,19^. Divergences during adolescent development of PFC GABAergic input have been proposed as a potential mechanism underlying the sexual dimorphism observed in psychiatric disorders, including anxiety^20,21,27^. Similarly, CB_1_ density, distribution, and binding properties differ significantly between males and females^22–26^. CB_1_ sexual dimorphism is also described after first-episode psychosis^28^ in humans, and in fear processing processes relevant for posttraumatic disorder in mice^29^. Taking all of this into account, CB_1_ signalling in GABAergic interneurons emerges as a possible pathophysiological player underlying emotional impairment in psychiatric disorders.

In this work, we have characterised the role of CB_1_ in GABAergic interneurons and mPFC function in both males and females. We described that pan-neuronal deletion of CB_1_ alters innate emotional behaviour, primarily in females, whereas loss of CB1 specifically from GABAergic cells primarily affects males. These behavioural deficits are linked to increased activity of these neurons.

Interestingly, this was inverted when we analysed learned emotional behavior using the fear-conditioning protocol. In this case, neuronal CB_1_ deletion increased the freezing response in males, while CB_1_ depletion in GABAergic neurons solely affected females. In the latter, these alterations were associated with reduced activity of GABAergic interneurons.

In summary, mPFC-GABAergic CB_1_ deletion induces an anxiolytic state and an exacerbated response to conditioned cues in a sex-dependent manner.

## Methods

### Animal models

*CB_1_*-flox animals were housed in the Animal Facility of the University of the Basque Country EHU. Animals were kept with water and food *ad libitum*, at a stable temperature and humidity. All animal procedures were approved by the Animal Research Ethical Committee of the University of País Vasco (M20/2019/199 and M20/2025/308 // M30/2019/301 and M20/2025/309).

### Surgery and virus administration

CB_1_ deletion was induced in 8-9-week-old *CB_1_*-flox mice, both male and female. Animals were first anaesthetized with isoflurane at 5% for induction and then maintained at 1% for the rest of the surgery. Breathing rate and body temperature were controlled during the procedure. Adenoviral vectors (AAV) carrying the Cre protein under different promoters were bilaterally injected at AP +2.0, ML±0.3, and DV −2.0, using the Nanoject III system (Drummond Scientific; Broomall, PA, USA). 200nl of AVV was injected at a 1nl/s per second rate. Lidocaine was topically administered at the beginning and end of the surgery, along with Meloxicam (2mg/kg) subcutaneously. Animals were kept in individual cages and monitored for 1 week.

For fiber photometry, an AAV encoding the calcium sensor GCaMP was injected simultaneously. After viral administration, a 3 mm fiber cannula (RWD; Guangdong, China) was carefully implanted at the same coordinates. The fiber cannula was secured using Superbond cement (Sunmedical, Shiga, Japan) and then dental cement (Methax Thermo, Makevale; Ware, United Kingdom) with carbon (Aquineuro; Bordeaux, France).

### Viral vectors

AAVs were acquired in the Viral Vector Facility VVF of the University of Zurich and used to induce CB_1_ deletion in the global mPFC neuron population (ssAAV-9/2-hSyn1-chl-EGFP_2A_iCre-WPRE-SV40p(A)) and in mPFC GABAergic neurons (ssAAV-1/2-mDlx-HBB-chI-iCre-WPRE-bGHp(A)). For tracking GABAergic neuron activity, we employed a GCamP8m (ssAAV-1/2-mDlx-HBB-chI-jGCaMP8m-WPRE-SV40p(A)).

### Behavioral Studies

Behavioural tests will start 5 weeks after surgery to ensure complete depletion of CB_1_ receptors^30^. Animals were habituated to human handling and to being connected to the fiber system for 3 days prior to the start of the behavioural tests. On the first day, animals were handled for 10 minutes. On the second and third days, animals were gently handled for 5 minutes, then spent 5 minutes in the home cage with the fiber connected. Innate emotional behaviour and cognitive tests were recorded and analysed with ANY-maze software (ANY-maze; Dublin, Ireland).

### Innate Emotional Behaviour

Anxiety-like behavior was tested using the open field and elevated plus maze, in this order. *Open Field*: an open-field arena (40 × 40 × 45 cm) was employed, divided into a “center zone” (aversive region) of 6 × 6 cm and the rest of the maze as the “safe zone”. Inside the safe zone, the “corner areas” of the maze were also studied (four areas of 1 x 1cm). Animals were placed in the center and allowed to explore freely for 5 minutes. *Elevated Plus Maz*e: the maze was divided into 4 arms, 2 open-aversive zones and 2 closed-safe zones of 30×5cm, situated at a height of 49cm. Animals were gently placed in the center and allowed to freely explore the maze for 5 minutes. In each maze, the time spent and the number of entries into the aversive zones, the time immobile in the safe zones, and the total distance travelled were analysed.

### Working memory

To assess working memory, we employed the Y-maze (6 x 30 cm, arm) that animals freely explored for 9 minutes. The number and percentage of spontaneous alternations, i.e., visiting the three arms in a row without repetitions, were analysed. The percentage of correct alternations was calculated using the *(number of spontaneous alternations/(number of entries to the A zone + Number of entries to the B zone + Number of entries to the C zone −2))*100* formula

### Recognition memory

The Novel Object Recognition Test was conducted to study recognition memory. For this test, the same open field arena was used. Habituation and acquisition were performed on the same day, with one hour’s separation. For the acquisition, we employed two equal grey balls at the centre of the maze. Twenty-four hours later, we performed the recognition test, changing one of the balls for a white square. All tests lasted 9 minutes.

To assess the memory impairment, we calculated the discrimination index (D.I.) using the (TN+TF): DI = [TN-TF]/[TN+TF] formula^31^. All objects were previously assessed so that animals would not show any preference.

### Learned Emotional Behaviour

In order to characterize complex emotional responses, we employed the Cued Fear Conditioning paradigm as previously described in Soria-Gómez E et al., 201^32^, using a Fear conditioning system (Imetronic; Marcheprime, France). Briefly, the acquisition was performed on day one, on a square grey Perspex chamber (25×25cm) and a grid floor. The animals were left to explore for one minute with white noise, followed by a tone (60 dB, 1.5 kHz) for 10 seconds, which co-terminated with a shock (0.7mA/1-s) during the last second. This trial was repeated another two times, with an intertrial interval of 20 seconds. Afterward, the animals were allowed another minute of free exploration. Twenty-four and forty-eight hours after the acquisition, the animals were presented with the same tone for three minutes without the shock to assess recall and memory fading. For these tests, the context was changed into a colored cylinder (25cm of diameter) and a rough floor. The animals were able to explore the arena for one minute before and after the test. The number of freeze-and-rearing cycles was normalised per minute.

### Fiber Photometry

GABAergic neurons’ activity was tracked during behaviour studies, using a R811/FR-11 Dual Color Multichannel Fiber Photometry System (RWD; Guangdong, China). To synchronise calcium recordings with behaviour events, we employed an ANY-maze Synchronisation interface (ANY-maze; Dublin, Ireland). Recordings were done at 100fps and 5ms of exposure, employing the 470nm laser at 100 mW and the 410nm laser at 26mW to normalise. Fiber photometry signal was analysed with the RWD software (RWD; Guangdong, China).

### Tissue processing

Twenty-four hours after the last test of fear conditioning, we proceeded to the mice’s euthanasia. Animals were deeply anaesthetized by an intraperitoneal administration of Ketamine/Xylazine (80/10 mg/kg body weight) and transcardially perfused with PBS (0.1M, pH 7.4) followed by 4% formaldehyde solution using a peristaltic pump (Perimax 12; Spetec GmbH, Erding, Germany). Brains were extracted and post-fixed in 4% formaldehyde for 24 hours and then included in 15% and 30% sucrose at 4°C for twenty-four hours each. Samples were stored at −80°C until use.

#### RNAscope

To analyse CB_1_ receptor depletion, an in situ hybridisation was performed at the Imaging Facility of Achucarro. Hs-CNR1-C3 (brain) - CB1 probe (591521-C3, Bio-Techne R&D Systems, s.l.u; Madrid, Spain) was used to detect CB1 mRNA, and the images were taken in a TCS STED CW SP8 super-resolution microscope (Leica; Wetzlar, Germany).

### Experimental Design and Statistical Analysis

For the statistical analysis, we used GraphPad 9 software (GraphPad Prism version 9.0.0 for Windows, GraphPad Software, Boston, Massachusetts, USA, www.graphpad.com). First, data normality was assessed using the Shapiro-Wilk test. For behaviour studies, an unpaired t-test with Welch’s correction was applied. If the data failed the normality test, a Mann-Whitney test was employed. Fiber photometry calcium recordings and fear conditioning studies were analysed using two-way ANOVA followed by Sidak’s multiple comparisons test. Area under the curve (A.U.C) values were analysed using the same statistical approach as the behavioural data. The results were considered significant when the p-value was < 0.05.

## Results

### Cell-specific deletion of CB_1_ in the mPFC

To validate our model, we quantified CB_1_ mRNA expression in the mPFC of control animals and animals with CB_1_ deletion in GABAergic interneurons of the mPFC (mPFC-i*CB_1_*-KO) (Fig. S1). A decrease in CB_1_ staining was observed in both male and female mPFC-i*CB_1_*-KO animals, compared to controls (Fig.S1). No signal was observed in the full *CB_1_*-KO animal (Fig. S1).

### mPFC pan-neuronal CB_1_ deletion modulates Emotional Response in a sex-dependent manner

The CB_1_ receptor is largely expressed in the mPFC, where it plays a key role in modulating neuronal activity^11^. To assess its role in behaviour, we selectively deleted CB_1_ expression in all neurons of the mPFC in both males and females (Fig. 1).

**Figure 1.**
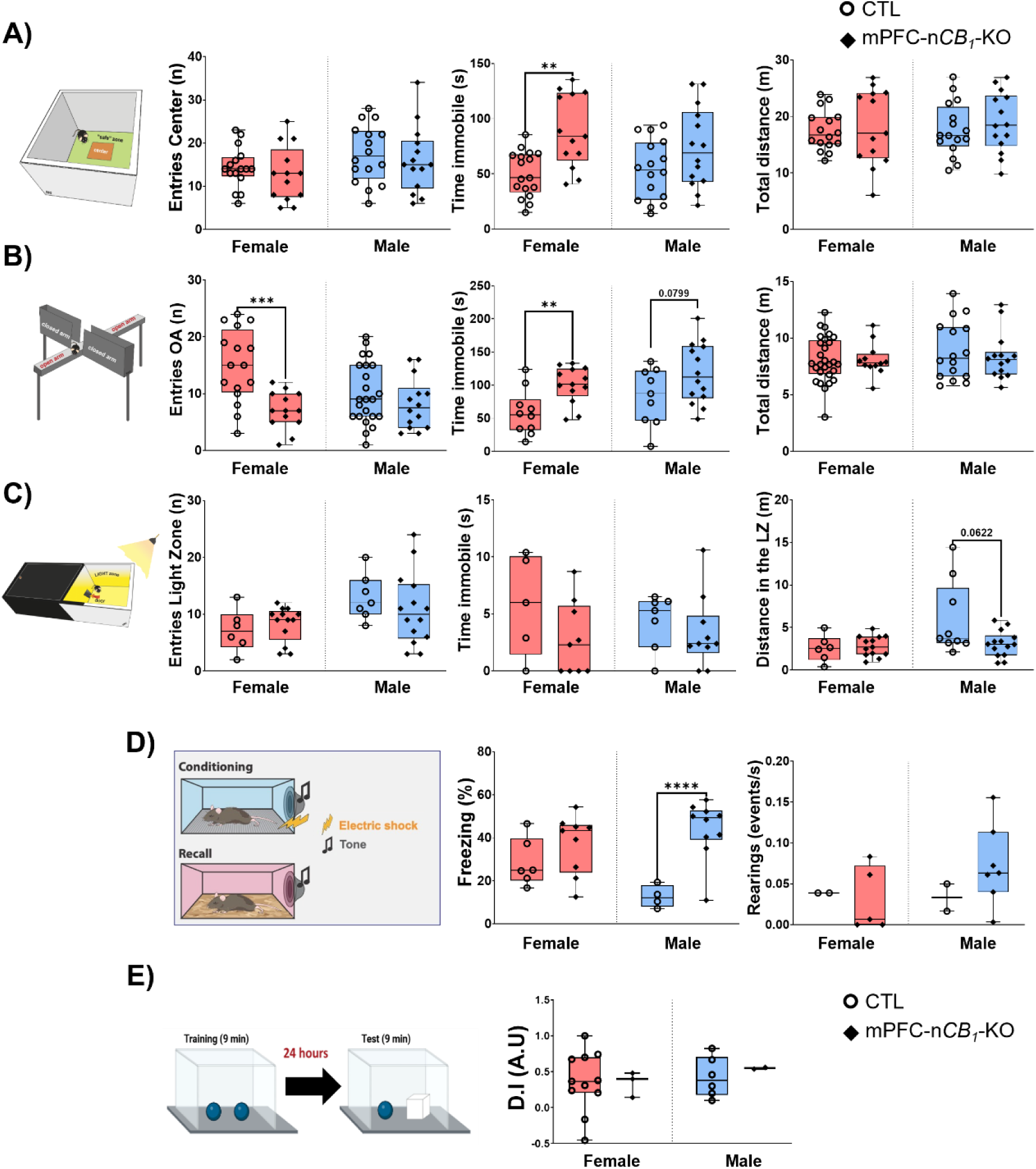
Neuronal CB_1_ deletion in the mPFC induced sex-dependent emotional behaviour changes. Five weeks after CB_1_ deletion, animals underwent different behavioural tests to assess their emotional and cognitive state. The innate emotional behaviour was studied in the open field **(A)**, elevated plus maze **(B)** and dark-like box **(C)**, in both female ***(red)*** and male ***(blue)*** animals. The number of entries in the aversive zone ***(left panel)***, the time immobile ***(centre panel),*** and the total distance travelled ***(right panel)*** were analysed in both controls (***circles***) and neuronal *CB_1_*-KO animals (***diamonds***). Complex learned emotional behaviour was studied through fear conditioning **(D)**, with an increase in the freezing ***(left graph)*** but not rearing response ***(right graph)*** detected. No impairment of the recognition memory was observed in the novel object recognition test **(E)**. Data are presented as median and the interquartile range, with min and max bars. Statistical analysis by Welch’s t-test. ** p<0.01; **** p<0.0001 vs CTL.

Pan-neuronal CB_1_ deletion primarily altered emotional behaviour rather than cognitive processes. However, its impact depended on the sex. The loss of neuronal CB_1_ had an anxiogenic effect in females, with increased immobility time in the open field and the elevated plus maze (Fig. 1A and B, center panel) and fewer entries into the open arms in the latter (Fig. 1B, left panel). The total distance travelled was similar between groups, which may rule out any motor impact induced by the deletion (Fig. 1A and B, right panel). No statistical difference was found in males, although there was a trend to an anxiogenic behaviour in most stressful mazes. Pan-neuronal CB_1_-KO male mice spent more time immobile in the elevated plus maze (p=0.079, Fig. 1B, center panel) and travelled shorter distances in the light box of the dark-light box maze (p=0.0622, Fig. 1C, right panel). In this line, the fear conditioning test showed that solely male mPFC-n*CB_1_*-KO mice had a potentiated freezing response 24 hours later, with no differences in the rearing frequency (Fig. 1D). In opposition to the emotional results, no cognitive impairment was observed in the novel object recognition test in female or male animals (Fig. 1E).

In conclusion, neuronal CB_1_ is a key regulator of mPFC-mediated emotional processing, but the mechanisms differ between sexes.

### CB_1_ receptor deletion in GABAergic neurons affects male innate emotional behavior

Since CB_1_ is mostly expressed in mPFC GABAergic neurons^8,11^, we next selectively ablated CB_1_ in this neuronal population (mPFC-i*CB_1_*-KO). In contrast with what we observed with global CB_1_ neuronal depletion (Fig 1), male, but not female, mPFC-i*CB_1_*-KO mice showed an impairment on the innate emotional behaviour (Fig. 2 and Fig. 3). In a less stressful test, the open field, the loss of GABAergic CB_1_ receptors decreased the time spent in the center (Fig. 2B) but did not increase the time in the “safest” spot, i.e. the corners of the field (Fig. 2D-E) neither the time immobile (Fig. 2F). Interestingly, GABAergic neurons activity was increased in mPFC-i*CB_1_*-KO female mice (Fig. 2G-H, left panel) when the animal crossed the center of the field despite no difference was observed in behavior (Fig. 2A-F). No difference in interneurons’ activity was found when the females moved to the safe zone (Fig. 2I-J), or in males in general (Fig. 2G-J).

**Figure 2.**
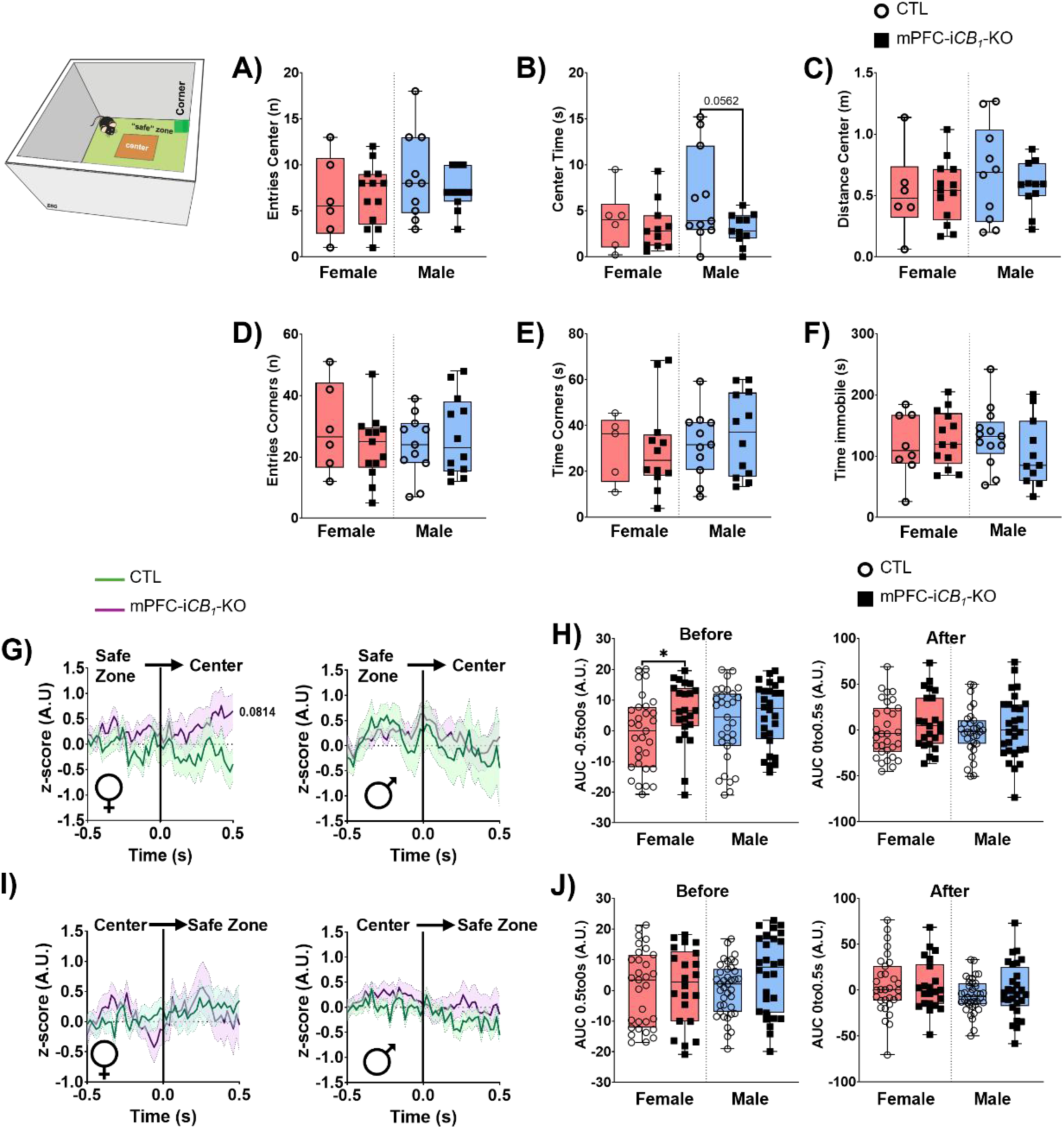
Effect of mPFC GABAergic CB_1_ deletion on cell activity and animals’ behaviour in the open field. Control *(CTL, circles)* and animals in which CB_1_ deletion was depleted in mPFC interneurons *(mPFC-iCB_1_-KO, squares)* were submitted to the open field test to assess the effect of CB_1_ deletion on animals’ innate emotional behaviour. The entries **(A)**, time **(B)** and distance **(C)** in the aversive center zone were characterised, with the only difference found in males’ *(blue)* center time **(B)**. Similar behaviour was observed in the number of entries **(D)** or time spent **(E)** in the safest zone, i.e. corners of the field. or in the time immobile **(F)**. GABAergic neuron activity during entry in the center **(G**, **H)** or in the safe zone **(I**, **J)** was studied. The curves of cellular activity (**G, I**) during the ±0.5 seconds of each entry showed no differences in female *(left panel)* or male *(right panel)* animals. A statistical increase of the area under the curve (**H**, **J**) was only observed in females (**red**) beforethe entry into the center (**H**, *left panel*) but not after (**H**, *left panel*). Box and whiskers data representation are presented as median and the interquartile range, with min and max bars. Statistical analysis by Welch’s t-test. GABAergic neuronal activity is represented as a mean ±SEM. Statistical analysis by a Two-Way ANOVA. * p<0.05vs CTL.

**Figure 3.**
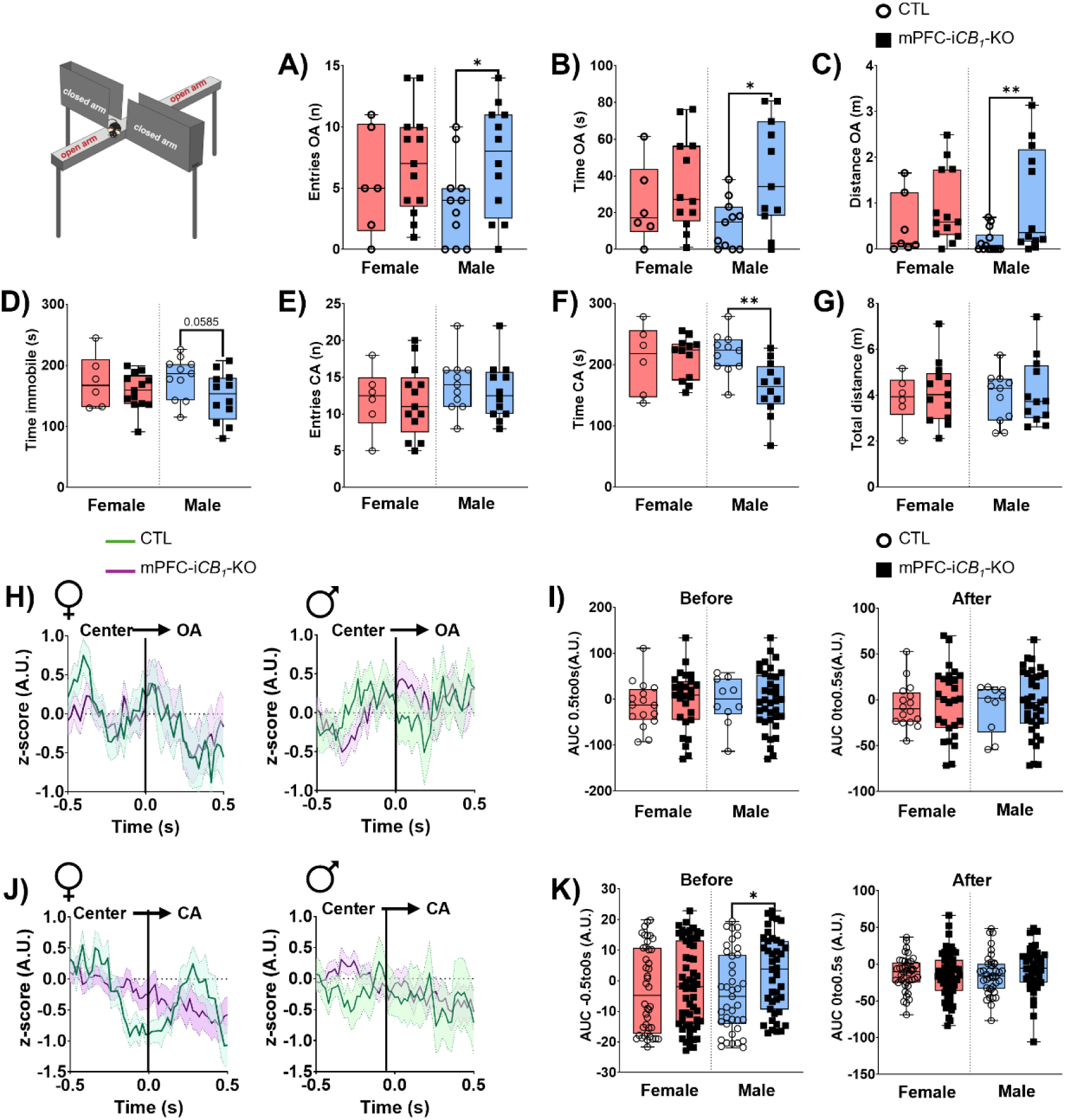
Interneurons CB_1_ deletion promotes anxiolytic behaviour associated with increased GABAergic neuronal activity in the elevated plus maze test. Innate emotional behaviour of both female *(red)* and male *(blue)* animals was studied in a more stressful maze: the elevated plus maze. A higher number of entries **(A)**, time spent **(B)** and distance travelled **(C)** in the aversive zone, i.e. the open arms *(OA)*, were observed in male animals with CB_1_ deletion in inhibitory neurons *(mPFC-iCB_1_-KO, squares)* versus control (*CTL, circles*). A trend to spend less time immobile **(D)** was also detected. The number of entries **(E)** or time **(F)** spent in the closed arms *(CA, the safe zone)* were only different in the latter. The total distance travelled was similar between animals **(G)**. While no statistical differences were observed between CTLs *(green)* and in mPFC-iCB_1_-KO *(purple)* concerning GABAergic neurons’ activity in the ±0.5 seconds of entering the OA or the CA (**H**, **J**). The studies of the area under the curve (*A.U.C,* I, K) showed an increase in males’ activity in the 0.5s before entering the CA (**K**, *right panel*). Box and whiskers data representation are presented as median and the interquartile range, with min and max bars. Statistical analysis by Welch’s t-test. GABAergic neuronal activity is represented as a mean ±SEM. Statistical analysis by a Two-Way ANOVA. * p<0.05vs CTL.

The innate emotional behaviour was then assessed in the elevated plus maze, a more stressful maze. In line with previous results, male but not female mPFC-i*CB_1_*-KO mice were affected by the receptor deletion (Fig. 3). However, while in the open field we described a decrease in the number of entries in the aversive zone, GABAergic CB_1_-KO induced an anxiolytic effect, with increased time, entries and distance travelled in the open arms (Fig. 3A-C) along with a reduction trend in time immobile (p=0.0585, Fig. 3D). These results correlated with a lower time spend in the closed arms (Fig. 3F). Similarly to the results of pan-neuronal deletion, no difference was found in the total distance travelled by the animals (Fig. 3G), which point out that the effects described were no motor related.

Remarkably, mPFC interneuron activity during entry into the open arms was equivalent between CB_1_-GABAergic KO and control animals in both sexes (Fig. 3H-I), despite the behaviour differences described in males (Fig. 3A-D). By contrast, increased activity of these neurons in male mPFC-i*CB_1_*-KO was observed in the 0.5s before entering the closed arms (Fig. 3J-K).

In summary, we conclude that CB_1_ deletion in GABAergic neurons provokes an anxiolytic phenotype in males, but not females.

### CB_1_ loss in mPFC GABAergic interneurons did not affect working and recognition memory

No cognitive impairment was observed after GABAergic deletion (Fig. 4), neither in working (Fig. 4A-C) nor recognition (Fig.4D-F) memory. In the Y-maze, there was no difference between control and mPFC-i*CB_1_*-KO animals in the number or percentage of correct spontaneous alternations (Fig. 4A-B), with similar distance travelled among groups (Fig. 4C).

**Figure 4.**
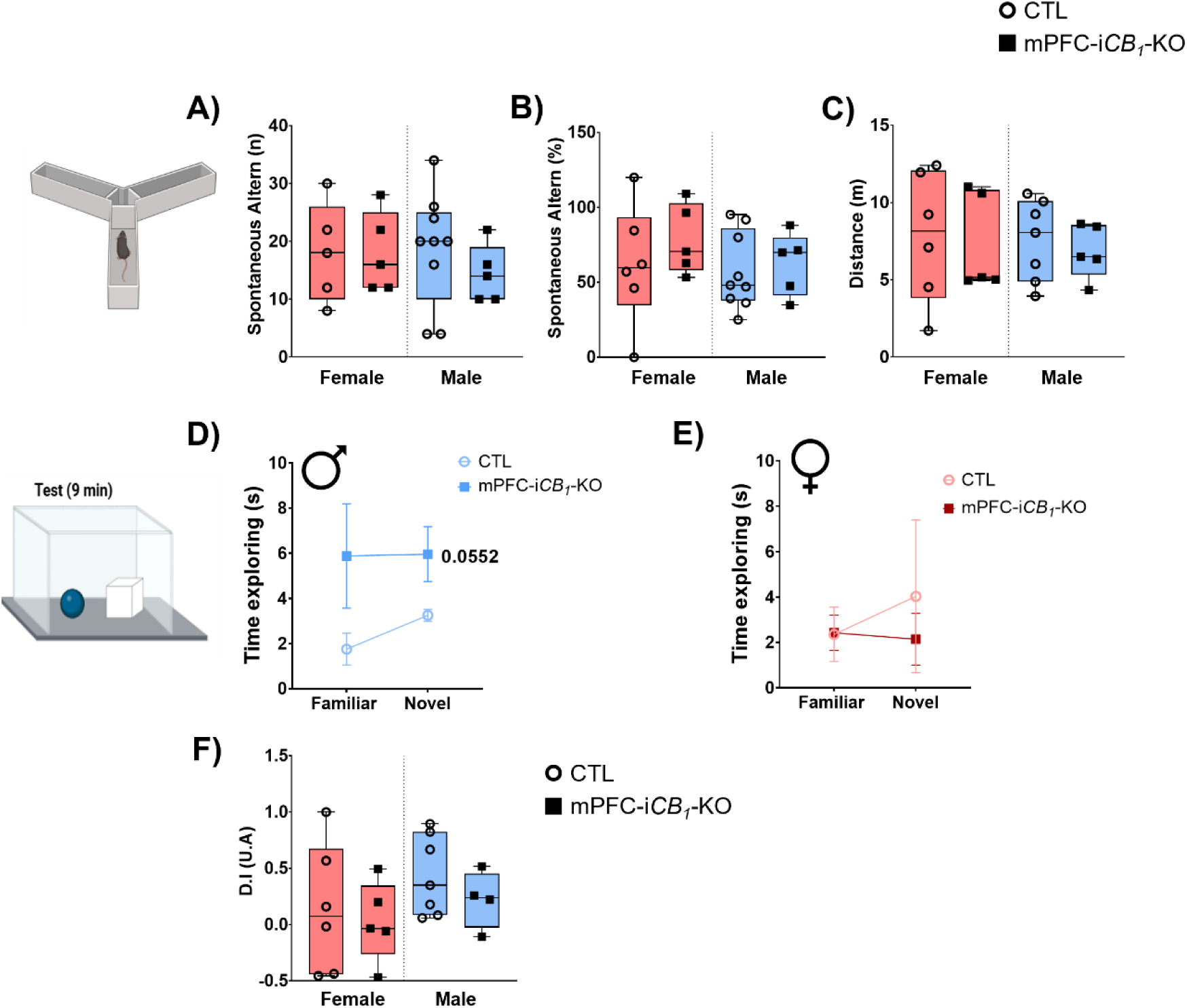
No main differences were detected on working or recognition memory after CB1 deletion in mPFC Interneurons. Cognitive impairment was studied using the Y-maze (**A-C**) and the novel object recognition (**D-F**) tests in control (**CTL**, ***circles***) and animals with CB1 deleted in inhibitory neurons of the mPFC ***(mPFC-iCB_1_-KO, squares).*** Both female (***red***) and male (***blue***) animals performed similarly in the Y-maze, regarding the number (**A**) or percentage (**B**) of spontaneous alternations and the total distance travelled (**C**). Similarly, there was no statistical difference in the discrimination index (D.I) of these animals in the novel object recognition test (**F**). However, a trend to spend equal time exploring both the familiar and novel object was identified in mPFC-i*CB_1_*-KO male (**D**) but not female (**E**) animals. Box and whiskers data representation is presented as median and the interquartile range, with min and max bars. Statistical analysis by Welch’s t-test. Time exploring is represented as a mean ±SEM. Statistical analysis by a Two-Way ANOVA.

To assess the recognition memory, we employed the novel object recognition test. While the D.I. was similar among groups (Fig. 4F), a clear increase in the exploration time was observed in males (Fig. 4D, p=0.0552) but not in female animals (Fig. 4E).

In conclusion, CB_1_ deletion from GABAergic neurons did not alter the working or recognition memory but increased the exploratory behaviour in males.

### Learned Emotional Behaviour was reinforced by CB_1_ depletion in the mPFC GABAergic interneurons of female mice

To assess learned emotional behaviour, we employed the fear conditioning paradigm, in which the mPFC plays a key role^33^. The association of the conditioned (shock) to the unconditioned stimulus (tone) was established on the first day (Fig. 5A), by employing three consecutive tones that ended with a shock, separated by an intertrial interval (ITI) of 20s. In general, no significant changes were observed in responses to the conditioned stimulus (Fig. 5B) or in animal activity during the ITI (Fig. 5C). However, there was a trend to an increased global freezing percentage in GABAergic *CB_1_*-KO females during the three ITI (p=0.0782, Fig. 5C, left panel).

**Figure 5.**
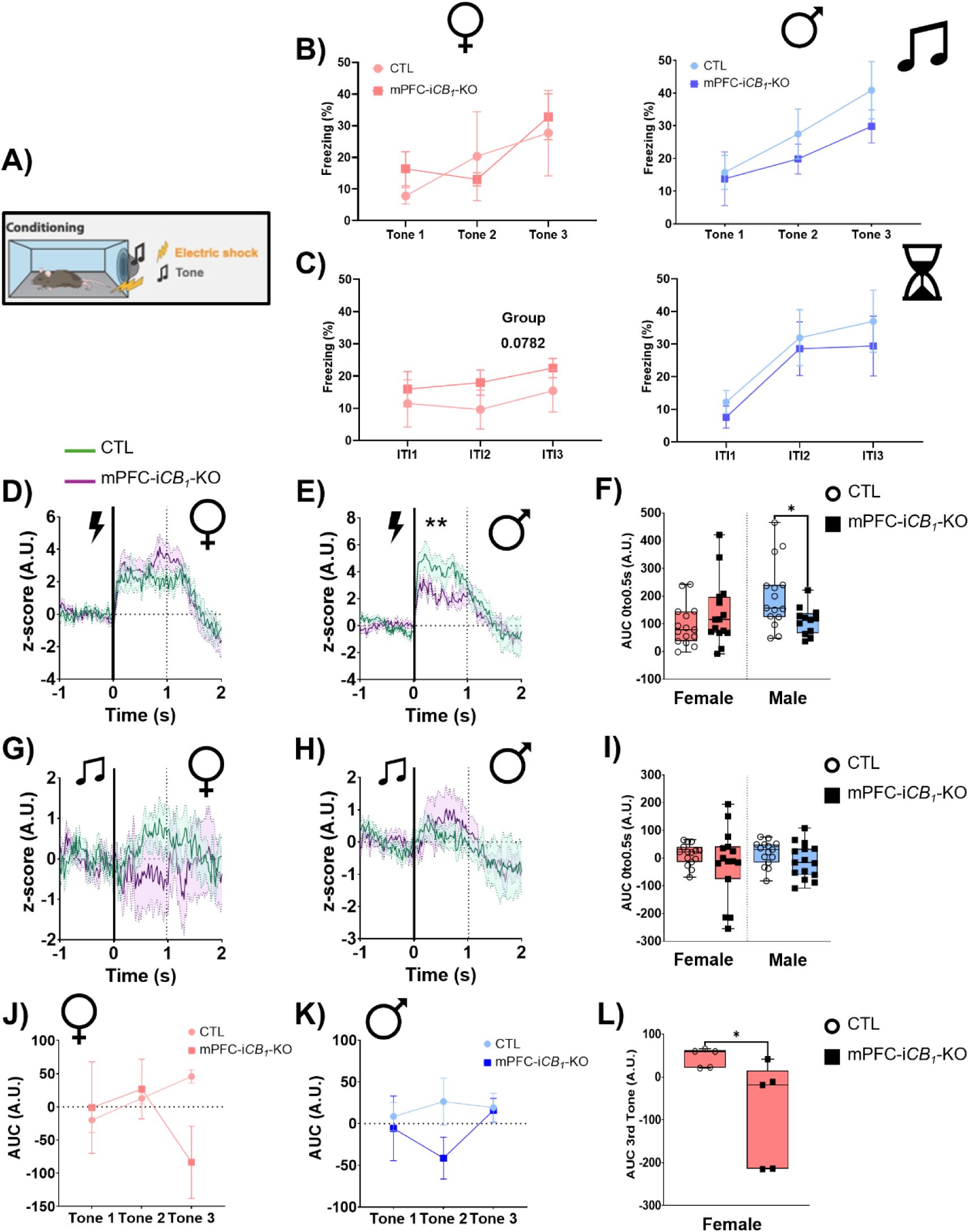
CB_1_ deletion in GABAergic interneurons of the mPFC potentiates freezing response during fear conditioning acquisition day, correlating with a decreased of inhibitory neurons activity. During the acquisition day, a tone was coupled to an electric foot shock three times with an intertrial interval of 20 seconds (**A**). The freezing response increased during the three tones (**B**) in both female (*left panel, pink*) and male (*right panel, blue*) mice without differences between controls *(CTL, circles)* and interneurons CB1-KO animals *(mPFC-iCB_1_-KO, squares)*. An increased freezing trend was observed in females during the three intertrial intervals (**C**). mPFC GABAergic neurons’ activity courses (**D-E**, **G-H**) from 1 second before to 2 seconds after the shock (**D-F**) or the tone (**G-I**) were also studied in both CTL (*green*) and mPFC-iCB_1_-KO (*purple*) animals. A significant difference between groups was found in GABAergic neurons’ activity in male animals (**E**), which correlated with a smaller area under the curve (*A.U.C,* F, I) in male animals (**F**). A.U.C. study during the three tones (**J**, **K**) was also studied, with a significant decrease in female neuronal GABAergic activity during the third tone (**L**). Freezing percentage and A.U.C during the tones and intertrials are represented as a mean ±SEM. GABAergic neuronal activity is represented similarly as a mean ±SEM. Statistical analysis by a Two-Way ANOVA. Box and whiskers data representation are presented as median and the interquartile range, with min and max bars. Statistical analysis by Welch’s t-test. * p<0.05; ** p<0.01 vs CTL.

While no statistical difference in female GABAergic neurons’ activity was found in response to the shock (Fig. 5D - F), there was a trend to lower activation after the tone (Fig. 5 G). Further analysis showed a steady increase of neuronal activity in control females, while in mPFC-iCB*_1_*-KO animals, said activity fell during the third tone (Fig. 5J). Indeed, interneurons’ calcium signal was significantly lower in mPFC-i*CB_1_*-KO females compared to controls (Fig. 5L). Along with this result, changes in the tone’s response during recall were only identified in KO females (Fig. 6). 24 hours after the acquisition, mPFC-iCB*_1_*-KO female mice showed an increase in the freezing response (Fig. 6D) linked to a significant decrease in the rearing response (Fig. 6E). This behaviour was not linked to differences in mPFC GABAergic activity (Fig. 6F). No difference in control and GABAergic CB_1_-KO’s responses were found 48 hours after the acquisition (Fig. 6B).

**Figure 6.**
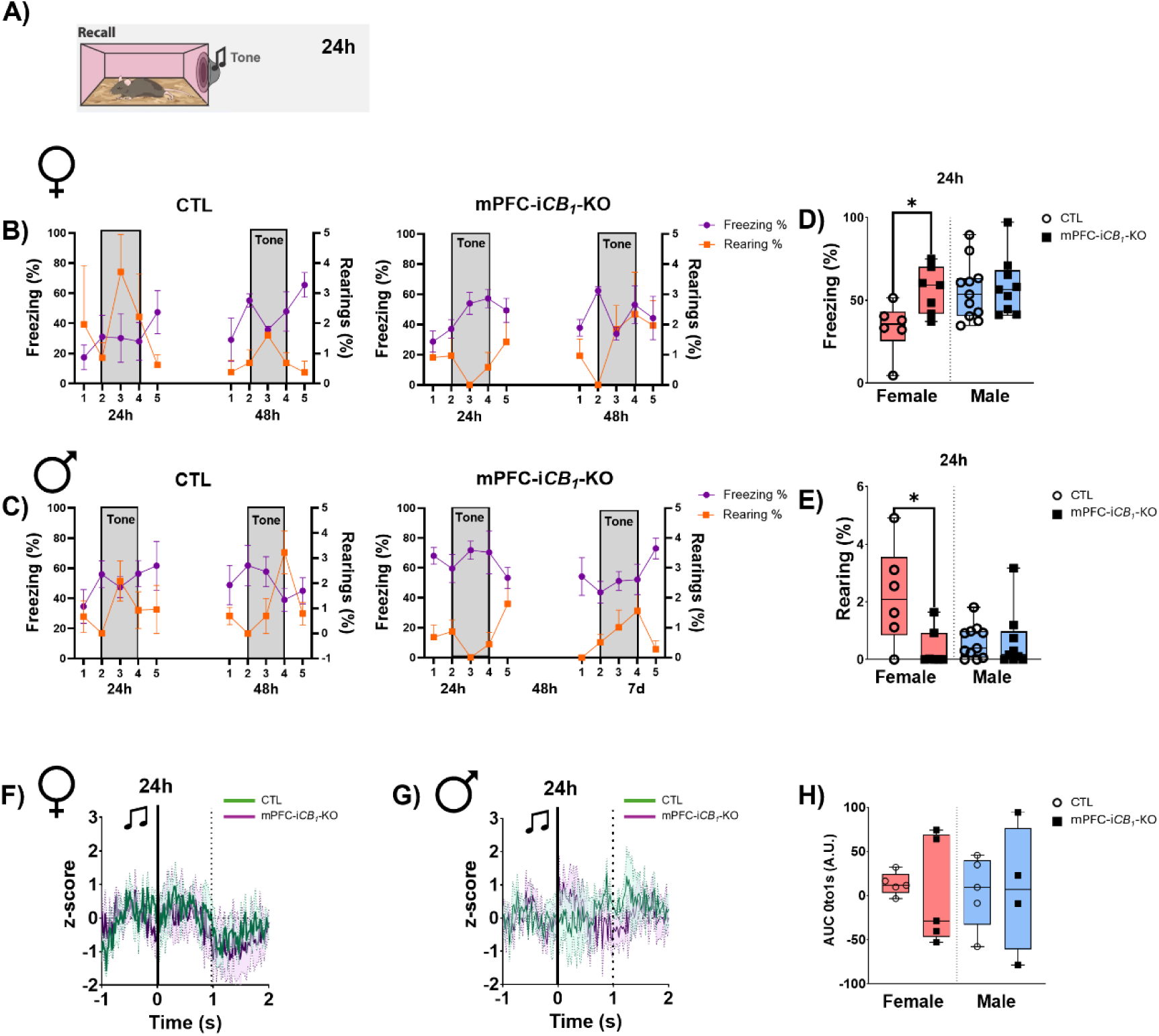
Females’ coping responses during fear-conditioning recall tests are altered after CB_1_ deletion in mPFC interneurons. Coping responses were assessed 24 and 48 hours after acquisition during recall tests, in which animals were exposed to the tone without foot shock (**A**). Freezing (***violet, circles***) and rearing (***orange, squares***) coping responses were measured during the 5-minute test period in both female (**B**) and male (**C**) mice, with an inverted trend reaction to the onset of the tone observed primarily in males (**C**). Significant differences were found in females (***red***) but not in males (***blue***) when the total percentage of freezing (**D**) and rearing (**E**) during the tone was compared. GABAergic neuron activity in the first two seconds of the tone was similar between control (***CTL***, ***green***) and animals without mPFC GABAergic CB_1_ (***mPFC-iCB_1_-KO, purple***) in both female (**F**) and male (**G**) mice. The area under the curve (AUC) of neuronal activity was also similar between groups (**H**). Freezing and rearing curves during recall tests are represented as a mean ±SEM. GABAergic neuronal activity is represented similarly as a mean ±SEM. Statistical analysis by a Two-Way ANOVA. Box and whiskers data representation is presented as median and the interquartile range, with min and max bars. Statistical analysis by Welch’s t-test. * p<0.05 vs CTL.

In male animals, we found an opposite pattern during acquisition (Fig. 5). While the freezing percentage was similar between controls and mPFC-i*CB_1_*-KO during the tones and the ITI (Fig. 5B and C, right panel), mPFC GABAergic activation was significantly smaller during the shocks in mPFC-i*CB_1_*-KO males (Fig. 5E-F). Regarding the tone, no differences were observed (Fig. 5H), although a different pattern of GABAergic neuron activation was identified during the three exposures, but with no significant differences (Fig. 5K).

Fear-conditioning recall was assessed 24 and 48 hours after acquisition, in which animals were presented with the tone without the foot shock (Fig. 6A). Freezing and rearing percentages were similar between control and mPFC-i*CB_1_*-KO male animals (Fig. 6D-E) 24 hours later, with no differences in GABAergic neurons’ activity (Fig. 6G-H). By contrast, further characterisation of the animals’ response did show some differences. While control animals reacted to the onset of the tone by increasing freezing and decreasing rearing, in animals without GABAergic CB_1_, the response was the opposite (Fig. 6C). This difference was observed not only at 24 but also at 48 hours after acquisition.

In conclusion, the loss of CB_1_ on GABAergic neurons alters fear-conditioning processing, potentiating the freezing response and decreasing the rearing activity, primarily in females.

## Discussion

The mPFC is one of the main regions involved in emotional processing, detecting potential conflict and participating in the expression and extinction of fear^33–36^. For this, the mPFC is connected with emotion-related areas like the basolateral amygdala, the mediodorsal thalamus or the hippocampus^5,7^. Inhibitory transmission in the mPFC plays a key role in integrating emotional and cognitive processes, maintaining an optimal excitation/inhibition balance^33,36,37^.

Given that CB_1_ receptors are predominantly expressed in GABAergic neurons within the mPFC^8,11^, we expected that pan-neuronal CB_1_ deletion, and specifically from GABAergic cells, would produce similar phenotypes. This convergence was observed only at the cognitive level, whereas the two manipulations had opposite effects on emotional behaviour. While pan-neuronal CB_1_ deletion selectively altered innate emotional behaviour in females, GABAergic-specific CB_1_ deletion affected basal emotional behaviour exclusively in males. A comparable, yet inverted, pattern also emerged during complex emotional processing. In the fear-conditioning paradigm, pan-neuronal CB_1_ deletion enhanced freezing responses in males, whereas GABAergic-specific CB_1_ deletion increased freezing behaviour in females.

These results point out 1) the relevance of the CB_1_ receptor on glutamatergic neurons in the mPFC, and 2) the clear sex-dimorphism of the endocannabinoid system in adult mice. Although in lower quantities, CB_1_ is also functionally expressed in mPFC pyramidal neurons, where it modulates glutamatergic synaptic transmission^38,39^. Previous studies have shown that CB_1_ deletion in cortical glutamatergic neurons alters basal animal activity and nesting^40^, and also modulates mechanical allodynia after nerve injury^38^.

Interestingly, CB_1_ expression in CaMKII-positive neurons is sex-dependent, with higher levels in females^38^. This would correlate with our results, in which the impact of global neuronal CB_1_ deletion on innate emotional behaviour was observed solely in this sex.

CB_1_ levels in glutamatergic neurons are decreased in pathological conditions, such as nerve injury^38^ or obesity, without altering GABAergic CB_1_^41^. This observation may explain our findings in the fear-conditioning paradigm, in which the enhanced freezing response was no longer observed in females but instead detected in males following neuronal CB_1_ deletion.

Concerning the role of this receptor in GABAergic neurons, our study showed that CB_1_ in mPFC interneurons is predominantly implicated in innate emotional behaviour in males, promoting an anxiolytic phenotype. Removing CB_1_ from GABAergic terminals presumably abolishes endocannabinoid-mediated disinhibition, thereby increasing GABAergic inhibition of mPFC pyramidal neurons and reducing prefrontal output. This correlated with the observed increase in GABAergic neurons’ activity in the open field and elevated plus maze. Our findings are consistent with pharmacological studies showing that direct enhancement of GABAergic tone in the mPFC via GABA_A_ receptor activation with muscimol, clonazepam, or GABA itself similarly produces anxiolytic-like effects in the elevated plus maze, whereas GABA_A_ receptor blockade with bicuculline is anxiogenic^42–44^. That aligns with the clinical use of GABA agonists for their anxiolytic effect^45,46^.

Interestingly, when we exposed the animal to a fear conditioning test, GABAergic CB_1_ deletion modified only females’ behaviour. Freezing responses generally increased during the acquisition phase, which may have contributed to the higher freezing observed during tone presentation 24 hours later. This interpretation is consistent with previous findings showing that stronger fear acquisition is associated with increased subsequent freezing^47^. This effect is directly related to the role of GABAergic CB_1_ in the mPFC since global GABAergic CB_1_ deletion reduces the freezing response^48^.

In this line, the infralimbic region of the mPFC plays a key role in the expression and suppression of conditioned fear^49^. Pharmacological or optogenetic activation of this area leads to a decrease in the freezing response during extinction trials^50,51^. In our study, deletion of CB_1_ receptors from mPFC GABAergic neurons resulted in a stronger freezing response, probably because of the presumed increased local inhibitory tone observed during the innate emotional behaviour.

By contrast, a detailed characterisation of GABAergic neurons’ activity during the acquisition day of fear conditioning showed a decrease in these neurons’ activation. Despite its controversy, other studies have shown that inhibiting parvalbumin-positive neurons, one of the main GABAergic interneurons of the mPFC, leads to an increased fear response^33^. Deeper studies characterising the effects of global GABAergic CB_1_ deletion on distinct interneuron populations are needed to understand the role of this receptor in complex emotion processing.

Finally, CB_1_ deletion in the mPFC did not impair working or recognition memory. The role of mPFC GABAergic neurons in cognitive tasks has been widely discussed, with controversial results^37,52,53^. A recent study has proposed that the involvement of mPFC GABAergic neurons depends on task difficulty, which may help explain the variability observed across studies^54^. Complementary experiments will be necessary to establish whether CB_1_ receptor function is similarly influenced by task demands.

To our knowledge, this is the first study to demonstrate the specific contribution of mPFC GABAergic CB_1_ receptors to emotional processing and to demonstrate that this role is sexually dimorphic. Global (GABA)*CB_1_*-KO, which cannot isolate the mPFC from the amygdala, hippocampus, or dorsal striatum, didn’t show any differences between males and females^55^. Moreover, the deletion per se did not affect the animal’s anxiety levels, with differences observed only after CB_1_ agonist administration^55^. By contrast, our study, focused on the mPFC, showed that not only were there basal differences in the innate emotional behaviour and coping responses to stress stimuli, but that these responses were sex- and cell-dependent. In this line, further studies are necessary to characterise the role of CB1 in specific subsets of GABAergic interneurons in emotional processing. Even though the dogma is that CB_1_ is not expressed in parvalbumin-positive neurons, our results pointed out that CB_1_ depletion in GABAergic neurons may modulate parvalbumin neurons’ activity, as described in the global *CB_1_*-KO^56^.

In conclusion, CB_1_ receptors present in GABAergic interneurons in the mPFC are key modulators of emotional responses in a sex-dependent manner. Thus, maladaptive prefrontal CB_1_ signalling could participate in the development of behavioural disorders.

## Acknowledgements

The authors would like to thank the Animal Facility and Arkaitz Pelaez (EHU), the Phenotyping Unit and the Imaging Facility of Achucarro for their technical and human support.

## Funding

This work was funded by the Spanish Ministry of Science and Innovation (PGC2018-093990-A-I00 and PID2021-125763NB-I00, funded by MCIN/AEI/10.13039/501100011033 to E.S.G.; PID2024-161059OB-I00 to S.M.); and cofounded by the European Union, the Basque Government (2023111031; IT1473-22, to S.M.; CannaMetHD, to S.M. and G.M.), M.C. was supported by the ExoPsyCogII grant.

---

**Figure S1.**
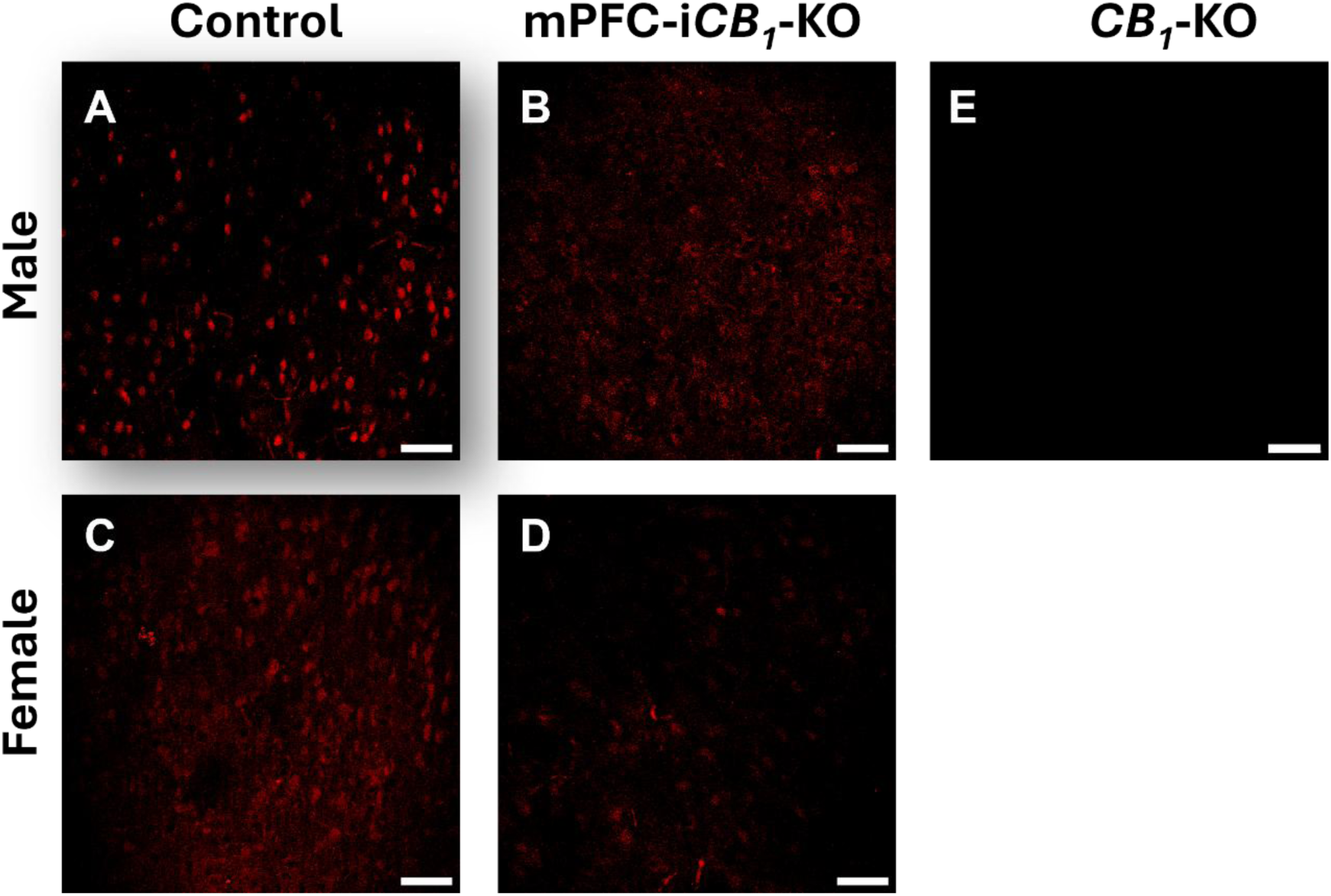
RNAscope of the CB_1_ gene in the mPFC of control and mPFC-iCB*_1_*-KO mice. Representative images of CB_1_ in situ hybridisation in the mPFC of male (**A-B**) and female (**C-D**) animals, which show a decrease in the cannabinoid receptor mRNA in mPFC-i*CB_1_*-KO animals (**B-D**) when compared to controls (**A-C**). Full *CB_1_*-KO animals (**E**) were used as a negative control, and no signal was observed in these animals. Scale bar, 20µm.

## Notes

### Competing Interest Statement

The authors have declared no competing interest.

